# Bird Species diversity from the Southern West part of West Bengal, India

**DOI:** 10.1101/2023.06.30.547191

**Authors:** Sourav Ch. Dinda, Arabinda Pal, Moulik Sarkar, Paresh Chandra Das, Niladri Mondal

**Affiliations:** Department of Zoology, Midnapore College, Midnapore,721101, West Bengal, India; Wildlife Institute of India, Dehradun, Uttarakhand 248001, India; Assistant Teacher, Chilkigarh I.C. Institution, Jhargram 721507, West Bengal, India; Kharagpur,721302, West Bengal, India; Department of Biology, KU Leuven, Leuven,3000, Belgium

**Keywords:** Checklist, Bird Diversity, Species Richness, West Bengal

## Abstract

Birds are essential for stabilizing the balance between various ecosystems. To estimate their diversity, we studied species distribution and made an annotated checklist along with their feeding habitats in the Southwest part of West Bengal, India. We studied the bird diversity using the point count method on various habitats, including agroforest, grassland, sub-tropical forest, and plateau forest regions. A total of 343 species were identified during our study period (2015-2019). We found high species diversity and sightings in the study sites, with 55 new records for the region. The whole study was conducted using a citizen science approach and amateur bird watchers’ community. This study will emphasize the future biodiversity conservation practice for the avifauna.

## Introduction

Human population growths establish and exploit limitless anthropogenic settlements, urbanization, and other development activities (Chalfoun and Schmidt 2012; Møller, Jokimäki et al. 2014). Due to extreme globalization throughout the world and the rise of the developmental phase is one of the most severe and irreversible forms of human impact, having far-reaching consequences for land use and land cover, hydrology and hydrobiology, associated biochemical activities, and climate change (Harris 2000; Baylis, Smith et al. 2017). It also significantly impacts monitoring natural resources connected with biodiversity, typically by converting natural or semi-natural habits, causing significant ecological changes worldwide (Dıaz, Symstad et al. 2003). Furthermore, almost every conserved tropical forest area is surrounded by human-occupied landscapes; thus, sharp focus on the avian biodiversity of esteemed regions to be partially or entirely affected by human activities (Eagles and McCool 2002). Unlike other higher vertebrate organisms, birds are crucial to the specific ecosystem and their corresponding natural habitat (Şekercioğlu, Daily et al. 2004). Therefore, most of the wildlife sanctuaries and national parks in India have been tagged with either one or more birds, stating them as flagship species in that largely conserved forest area.

Remarkably, fifteen species of birds in India were declared critically endangered by the International Union for Conversation of Nature (IUCN) in 2013. The endangered birds, including the Great Indian Bustard, Siberian Crane, White-backed Vulture, and Red-headed Vulture, are on a declining trend shown by the report of IUCN updated till December 2013.

The primary reasons for such a decline in the population of these avian species included loss, modification, fragmentation, and degradation of natural habitat, environmental contaminants, poaching, and land use changes, particularly the conversion of large areas for crop cultivation. Also, changes in cropping patterns are linked to various reasons, including implementation of irrigation schemes, increased pesticide usage, livestock-grazing, high levels of disturbance, and developmental activities like mining and hydel projects (Şekercioğlu, Daily et al. 2004).

American Museum of natural history states that nearly 18000 extant bird species are now available throughout the world (Richa Singh 2018). Among them, 1263 bird species have been identified in India by previous researchers (Praveen et al.). This comprises 12% of total bird species worldwide, while India enjoys only 2.4% of the global land mass.

Earlier studies have been done on avian biodiversity in the West Bengal region following the number of years gap (Sivakumar, Varghese et al. 2006). More than 949 bird species were recorded from India’s entire West Bengal province. Proper research has been done about the bird diversity and distribution in the northern region of West Bengal (Datta 2011) (Roy, Pal et al. 2011), but the least quantity of studies have been done on the western region of the specified state (Patra and Chakrabarti 2014).

Furthermore, situated in the western part of the West Bengal state, India, Diverse natural habitats, and ample sources of wetlands enable the specified region for various bird habitats. In this research design, a booming drive is also taken to prepare the latest avian species checklist, their conservation status, and habitat distribution patterns. A sharp approach leads to the methodological analysis of available avian species and diversity status specific to the Paschim Medinipur and Jhargram regions for estimating the variation of localized species richness and abundances.

This study aims to inquire about the bird diversity in Paschim Medinipur and Jhargram district, which will be conducive to preparing baseline data on bird diversity. This type of work is inevitable for the ornithological workers of these areas to reinvestigate the status of natural habitats in the future.

## Materials and Methods

### Study area

Paschim Medinipur and Jhargram districts are two adjoining districts located in the southwestern part of West Bengal. It is spread over a 9,345.64 km2 area (fig 1). This vast area includes forest, dry plateau, agricultural land, riverbed, and marsh wetland. These diverse habitats create a Suitable place for both resident and migratory birds. In connection with the Chota-Nagpur Plateau in the western part, rich faunal diversity is prevalent in that particular region (Mistry 2015). A large area of Paschim Medinipur and Jhargram is covered with Shal Forest, and few numbers of species prefer this forest for nesting. Four main river Subarnarekha, Kangsabati, Rupnarayan and Shilabati. Jhargram District and the western part of the Paschim Medinipur District are part of the Chotonagpur plateau (fig 2). This plateau consists of a dry hilly area and the central portion of the Shal Forest. During the summers, temperatures are high in this region, most vegetation is dried up, and most birds choose the shal (*Shorea robusta)* jungle as a shelter (except aquatic birds). Still, it is the rainy season with a mass vegetation growth, and it lasts until winter, which helped to get the bird diversity here. Agriculture, especially the vegetable fields, is a valuable attraction for Cuckoos, Shrikes, and other small predator birds for food availability during monsoon and winter.

**Figure 1.**
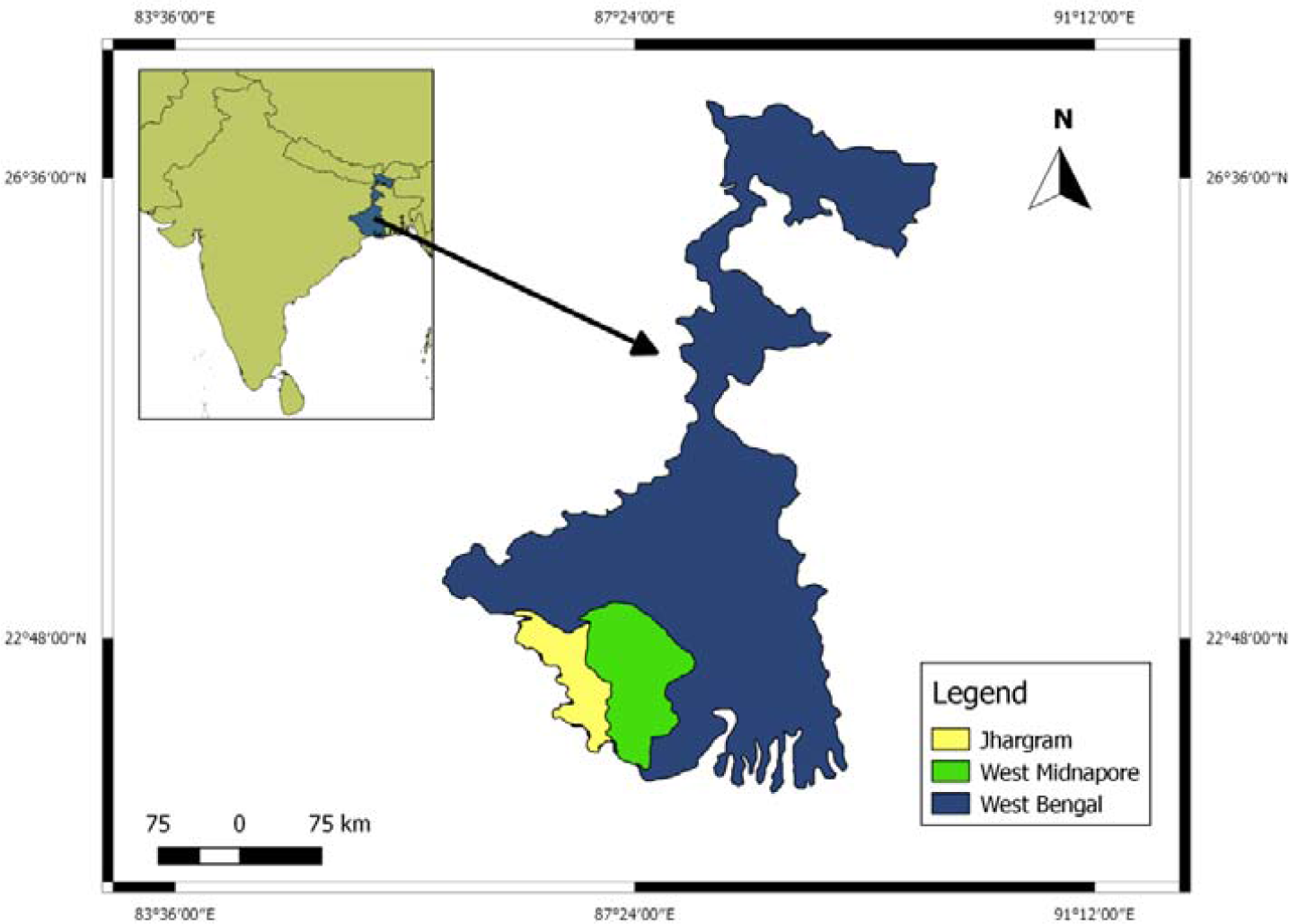
Preliminary survey site.

**Figure 2.**
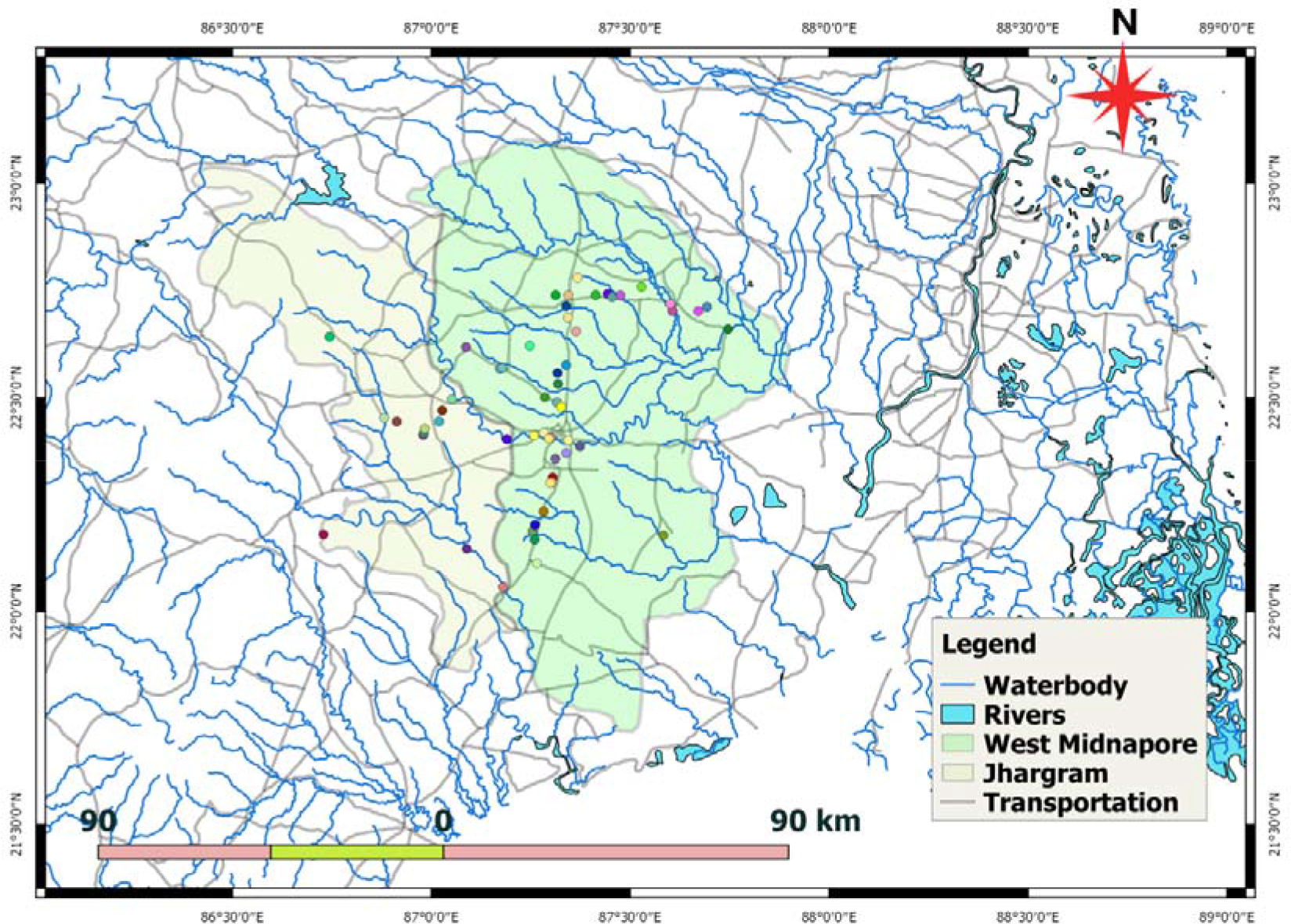
Map showing the study areas at Paschim Medinipur and Jhargram, West Bengal, India (including the water-body network)

#### Instruments used

The tools used was Zeiss conquest HD binocular 10x42 for watching and tracking and Nikon 60x digital zoom camera, Canon 40x digital zoom camera, Nikon D7000 with 150-600mm lens, Nikon D3300 equipped with a Nikkor 55-200mm lens for photography and closed observation. For proper identification of specific birds up to species level, typical guide manuals like the Birds of Indian Subcontinent by Richard Grimmet (Grimmett, Inskipp et al. 2013) and The Book of Indian Birds by Salim Ali (Ali 1943) were used, and specific morphological features also verified.

#### Survey technique

Recognizing an avian species can be a challenging scheme. Birds are diligently active and energetic creatures, so we need to spot them briefly or quickly. An unswerving survey was done depending on their calls, movements, feeding habits, and locomotive ground. A time gap field survey in regular mode was also carried out on those two Districts for four years, from July 2015 to July 2019, to study the avifaunal diversity. Every point has been sampled multiple times yearly, mainly based on birds’ perching times and accurate breeding times. For better bird sample count, sun rising moments and the areas closer to water bodies were randomly chosen in whole study periods (Ralph, Geupel et al. 1993). Line transects and point transects methods were used for this study; in this method, a substantial amount of traveling is required using a linear point which can be estimated by the naked eye, following survey estimation and sampling of bird species on both sides. This method is faintly similar to the point transect method (Gregory, Gibbons et al. 2004).

Ten sites are randomly selected and considered more species-rich zones to investigate the avifaunal real-time distribution. The desired places are chosen based on their vegetation status, practical food availability and bio-resources for birds. Targeted businesses, including the type of habitat complexes, are showing avian species variability findings in random mode (Table 1).

**Table 1:** Habitat types of the study areas (specific code name) for bird diversity.

| SL. No | Site name | Site code name | Habitat complex |
| --- | --- | --- | --- |
| 1 | Cossye riverbed and adjacent areas | CRA | Riverine segments, Shurbs, Sub-tropical forest |
| 2 | Khargapur And adjacent forest areas | KGP | Sal forest (sub-deciduous) |
| 3 | Barkola wetland | BWL | Grassland, Wetland , Farmland |
| 4 | Vosraghat and adjacent areas | VAA | Grassland, Farmland, Cultivation land |
| 5 | Teak forest zone of Arabari, shalboni, lalgorh | ARA | Dense Teak forest (deciduous) |
| 6 | Ghatal and adjacent area | GHA | Grassland and water washed land |
| 7 | Khargapur and CK road plateau | CKR | Grassland, sal forest (sub deciduous) |
| 8 | Jhargram Teak forest area | JHA | Teak forest (deciduous) |
| 9 | Jhargram Dam areas and wetlands | JDA | Water body, wetlands |
| 10 | Belpahari and adjacent areas | BEL | Deciduous forest , plateau |

**Table 2:**
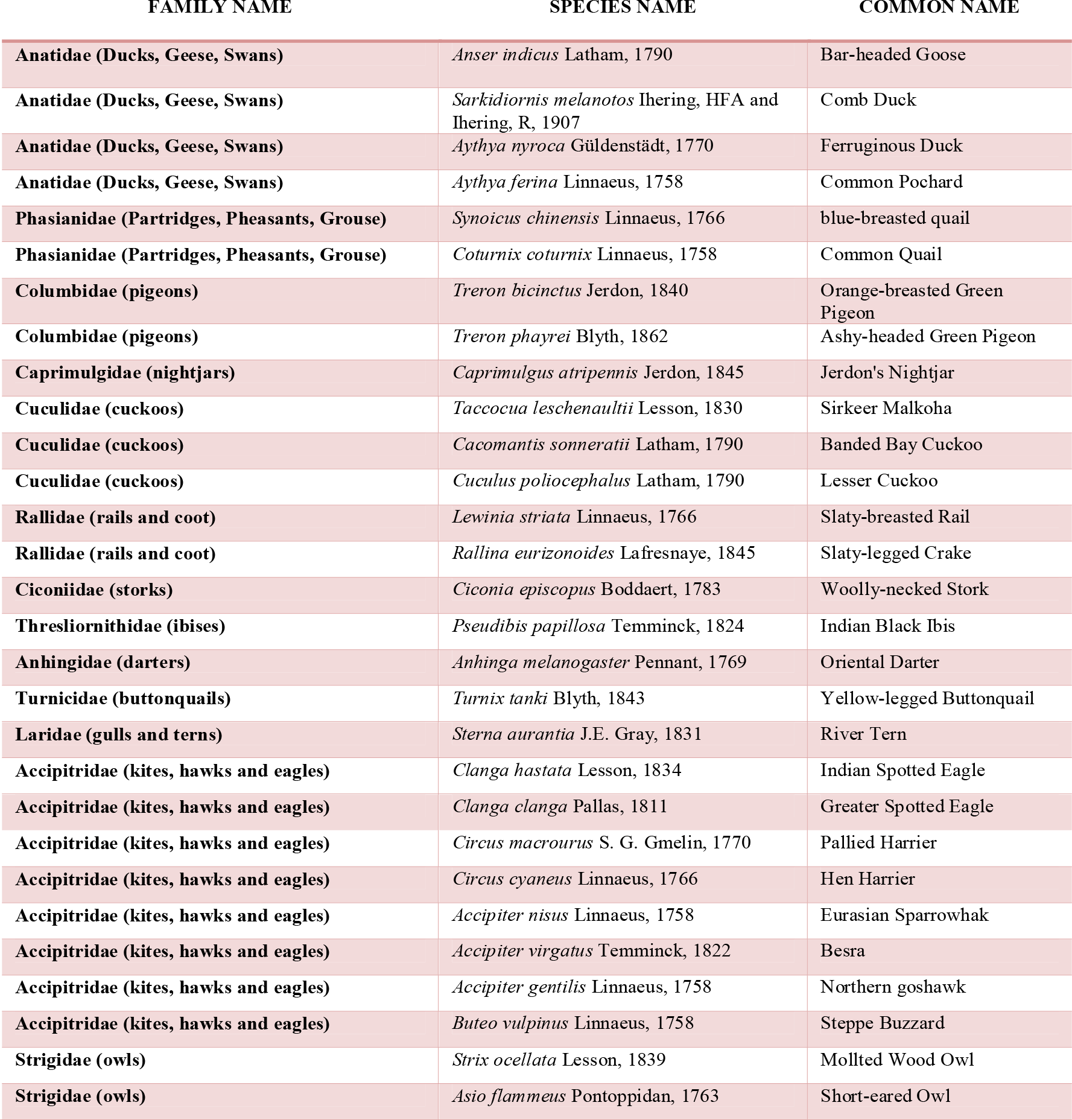
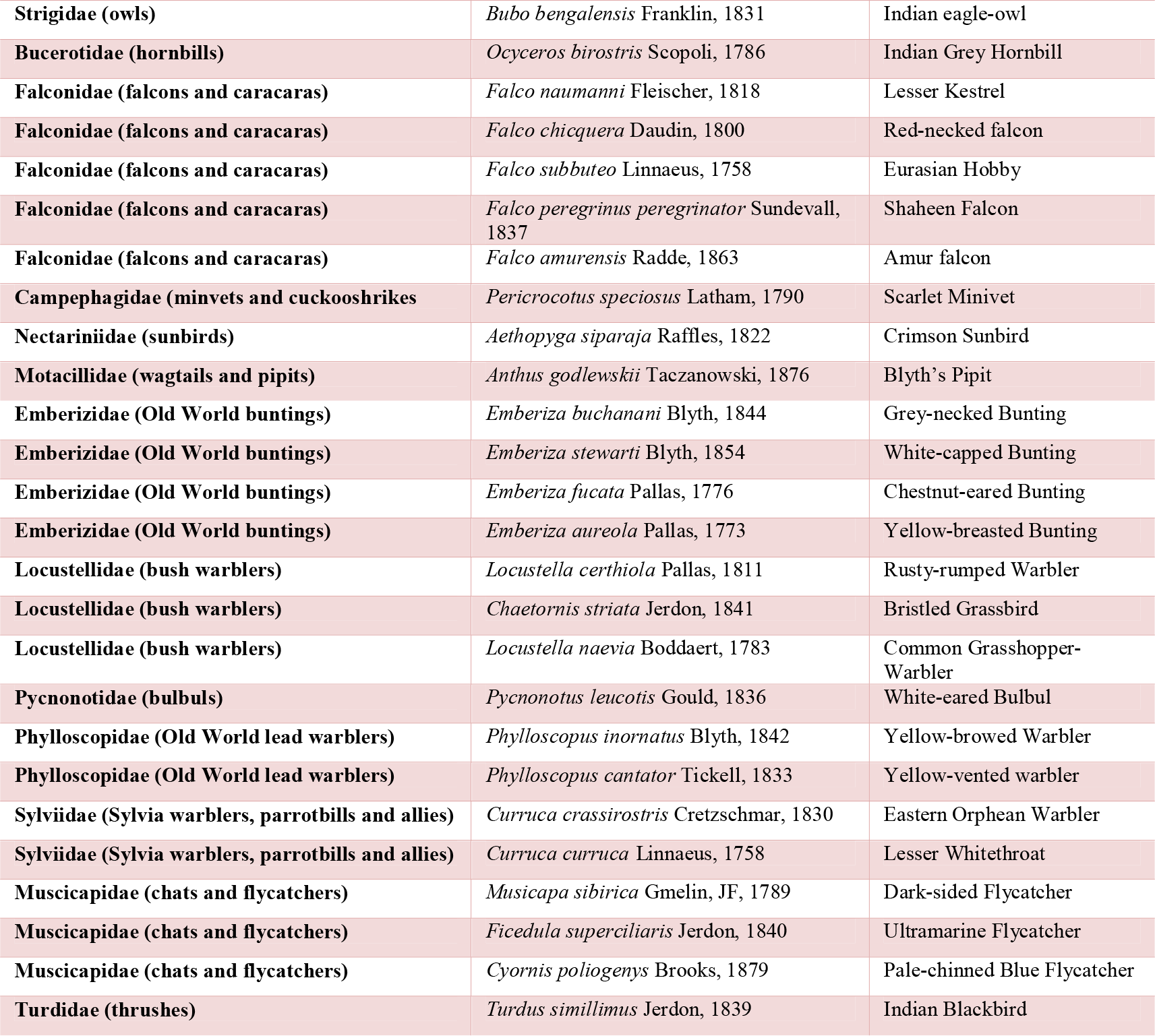
Checklist of New record from the study site.

### Habitat

These bird species have been found mainly on a) deciduous, shal-rich forests, b) wetlands, c) riversides, d) large water bodies, and e) human settlements. Due to the availability of diversified significant natural habitats and the abundance of forest areas, avian faunal diversity is well-reserved in different parts of Paschim Medinipur.

### Abundance of faunal diversity

Most bird species are prevalent within the wild habitat, but fewer are reported near or in the human settlement areas. Region-specific abundances of large water bodies and flowing river numbers of birds having migratory visit patterns like Rudy Shelduck, Common coot, and Lesser Whistling Ducks are spotted in the water bodies during the winter season. The measurable avifaunal diversity in the focused region has been calculated by using the Shannon-Weaver Index for estimating the species richness (Longuet-Higgins 1971).

### Species richness

In observatory periods, research also established that every bird species was visualized or heard back using the call method to consider bird abundance. Scientific calculation of species richness is one of the essential parts of conserving and measuring species biodiversity. At the same time, numerous biologists exploit individual evaluation models for investigating a few unique species at random manners and then identifying the desired species. Another functional technique supports enlisting all the individuals at once and ideally identifying the bird species individually. Such methods are based on sample assessment performance. These sample-based and individual-based species lists are calculated through statistical analysis (Pino, Rodà et al. 2000). In consider the non-parametric approach, which is more fluidic and overcomes the boundaries to detect the bird species abundance data. During the complete analysis schedule of the nonparametric distribution model, three main estimators are considered good standards throughout the perspective of the whole world.

### Statistical Analysis

Samples were tested in various statistical analyses using Past software to evaluate the collected data properly.

### Chao1-estimator

This estimator is based on species abundance. As the data is referred to the plenty of individuals, the data set concerns the abundance ratio of species individuals grouping in a particular class of samples. As so many species are present, they are only represented by one or two individuals citing them as rare species.

The Chao1 estimator emphasized this rare species accumulation. This type of estimator is based on lower-order species occurrence consisting of singleton and doubleton abundance data (Gotelli and Chao 2013).

To calculate the Chao1 estimator:, this is the bias-corrected method.

Where:

S_obs_= the number of species that occurred in a set of samples.

*f*_*1*_= Number of singleton species found in a sample

*f*_*2*_= Number of doubleton species found in a sample

### Species diversity

Species diversity is the quantitative estimation of species in a sample. To estimate the diversity, we have used the Shannon index and Simpson 1-D index method (Keylock 2005).

### Shannon index

It is known as the Shannon-diversity index or Shannon entropy. This method is based on the degree of ambiguity and prediction (Allen, Kon et al. 2009).
n= Number of individuals

### Simpson index

This index has been used to classify the degree of attention into types. This method took datasets of randomly placed samples of different kinds of samples (Jost 2006).

### Individual rarefaction analysis

To estimate the diversity at the taxonomic level, samples are added from various sites, sizes, and numbers. The abundance data can be calculated if the sampling is done from similar habitats using the standard method. The rarefaction data analysis method, the model estimate number of taxa found in a sample, and the comparison of different data sizes are also evaluated here. The log Gamma function using the computational method, the standardized graph, has been plotted (Krebs 1989).
Here, N= The number of individuals in a sample taken

S= Total number of species in a model, N_i_ = Number of individuals of species (number i)

E(Tusnady and Simon)= Expected frequency of species in a sample (n)

### SHE analysis

SHE analysis interprets the relationship between Species Diversity, Richness, and Species Evenness index. In SHE analysis, the number of species and equitability are examined, and the parameter depends on the sampling methods used during observation. Here, S is the species number in a sample; H denotes the information fraction; E is the model’s evenness measure. If abundant species occurrence data is present, SHE analysis can compute the overall community structure of the sample area.

## Result

Here we got the first annotated checklist of birds for the Paschim Medinipur and Jhargram district of West Bengal, with 343 species in total with their sightings and their habitat. We got bird species from 69 different families, which clearly indicates the huge diversity around the study area (supp Figure.1). This study was crucial to assess the conservation status as well as to look more into how many species are resident or migratory in the region (Figure.3).

**Figure 3.**
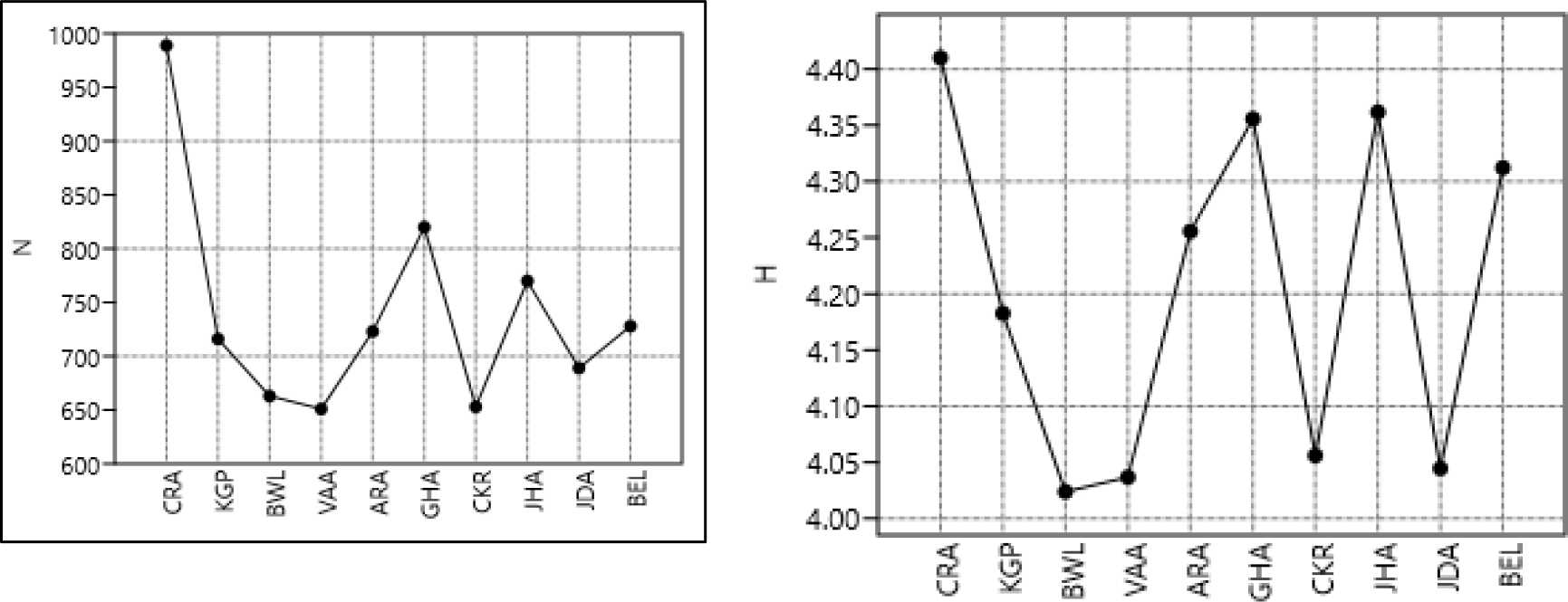
Chao-1 estimator and Shannon index (entropy) for study sites.

**Figure 4.**
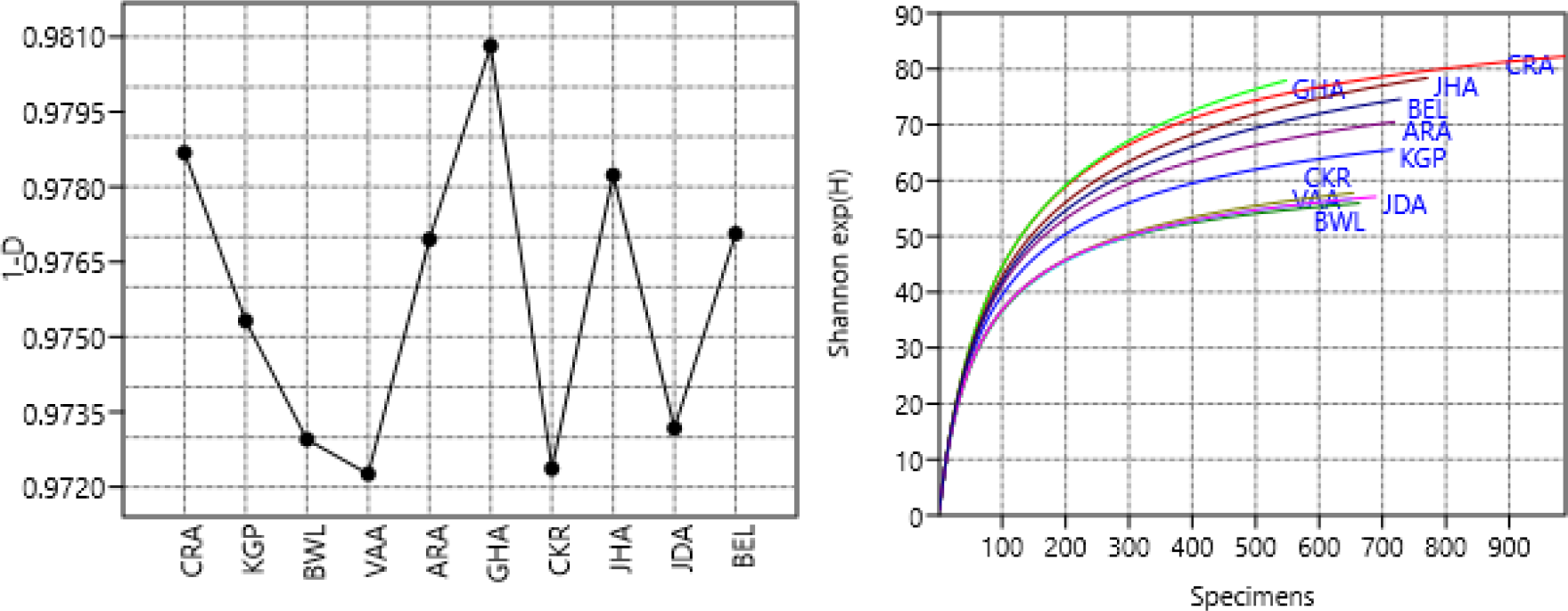
Simpson’s (1-D) index and Individual rarefaction analysis for the study sites.

**Figure 5:**
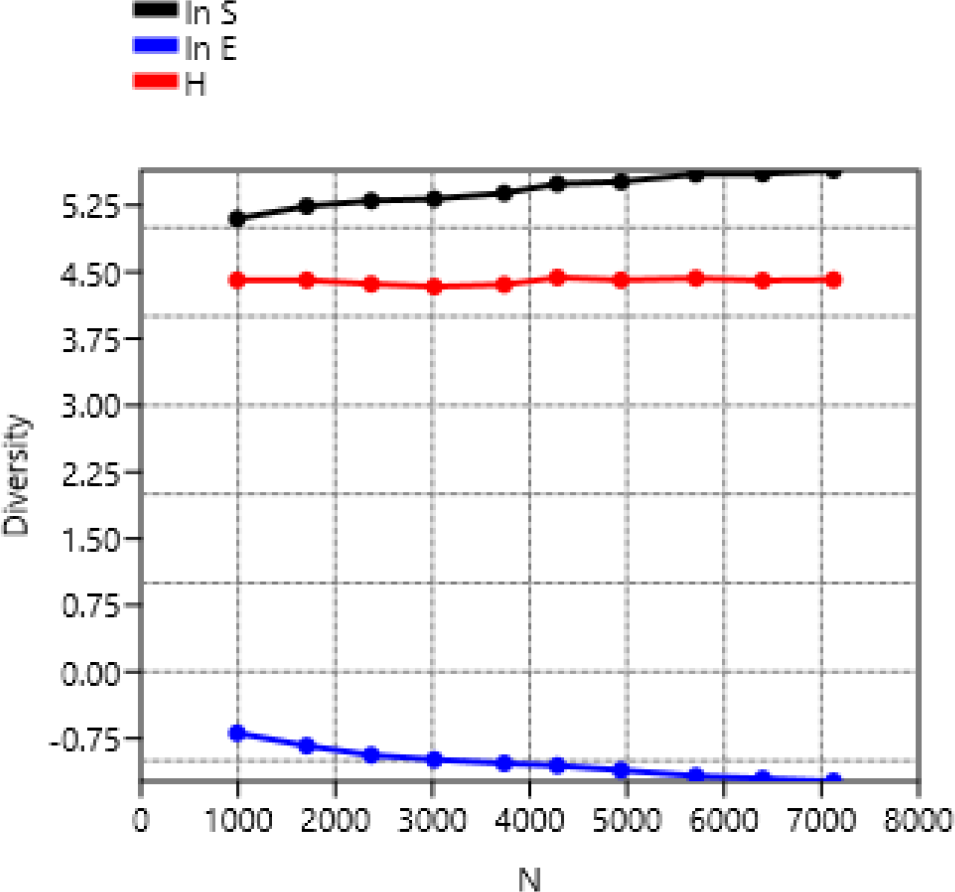
SHE analysis for study sites.

### Bird species list recorded: Checklist

Three hundred forty-three bird species were recorded from 2015 to 2019 from the six sites (supplementary file). This is the first annotated checklist from the southwest of West Bengal, with intense surveying and covering almost all types of habitats.

**Fig. 3a.**
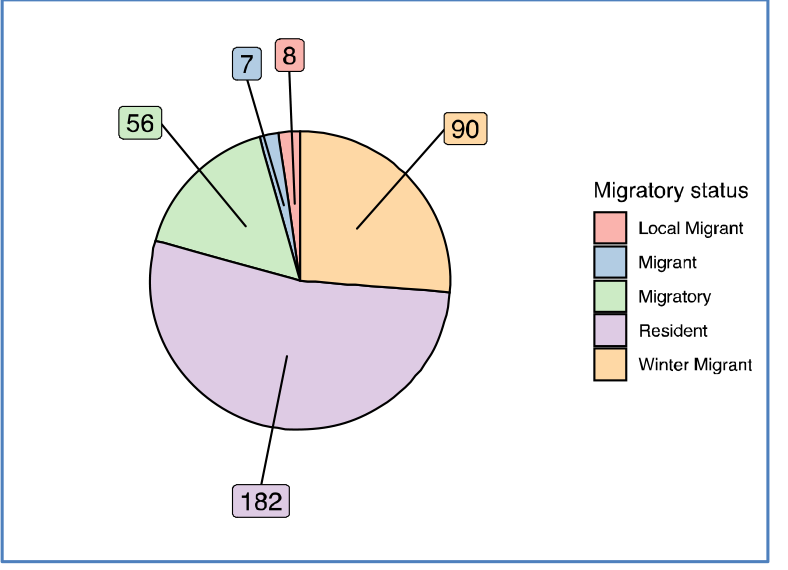
Migratory Status.

**Fig. 3b.**
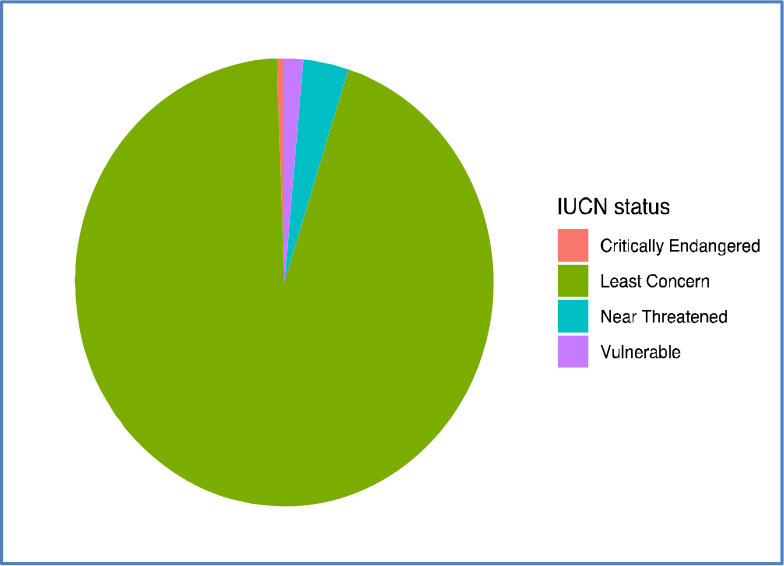
Conservation status.

### Species diversity and Species richness

Calculated estimators (Chao_1, Shannon entropy) show that sample sites CRA, GHA and JHA are more species-rich than the remaining seven spots. This is due to the river ecosystem, which acts as a feeder for the bird community. (Hughes, Daily et al. 2002)

The index shows that more individuals are present in the Ghatal and adjacent areas reflecting on the established index. The high abundance of individuals in places CRA and JHA also reflects on the graph due to their high species richness and occurrence of maximum individuals due to abundant water and food sources in those areas.

### Individual rarefaction analysis Result

From the rarefaction analysis of the data taken from the sample sites. Site CRA has the maximum number of data variance as well as species diversity. The rest of the sites are then periodically following the same order. Site JHA and GHA are similar in species diversity and abundance ratio about taxonomic similarity.

From the above graphical data, the ln E value and ln E values are calculated. The H value has been constant due to the synchronised decreasing and increasing order value of the other two data (lnE, ln S). The value of H is reduced very slowly as the overall species abundance rate is spread n a significantly vast range. From the data, general community structures of the sampling sites are revealed, and extensive species occurrence and favorable species habitat are inferred.

## Discussion

Statistical analysis shows that the three distinct areas (GHA, CRA, JHA) are more diverse and have a high turn up of species due to the presence of forests, wetlands, rivers, and an enormous amount of food sources; the remaining seven areas have a low amount of bird diversity as this areas are not diverse as the three areas(GHA, CRA, JHA). The ten sites recorded three hundred forty-three bird species from July 2015 to July 2019. Concerning individual species counting and field data survey, areas adjacent to large water body employs a high number. Cossey River contains water bodies, grassland, forest, and cultivated land, making this region more suitable for rearing different birds. Dense forests with undergrowth support small birds (family) for other purposes, viz. nesting material, lower predation rate, and undisturbed breeding ground. They result in the availability of Indian pitta, flycatchers, cuckoos, thrush, and other birds.

## Conclusion

Habitat fragmentation is the most common threat these species encounter throughout the year. Construction of roads and human settlements play pivotal roles in destroying the natural habitat of avifaunal species (Rajashekara and Venkatesha 2017). Threats to avian diversity are the same on these field sites. The variety around the Kasai-Halt is gradually decreasing due to constant human presence as a disturbance to the ecosystem. Recent activities like the Brick-Kiln industry (Skinder, Sheikh et al. 2014), road construction, and heavy vehicles on the riverbank for the sand mine industry hamper the ecological balance. We have recorded a gradual decrease of species like Little Ring Plover, Sandpiper, and White Wagtails in the riverbank. Status of Gopegarh, although it is a reserve ecological park, habitat fragmentation has also been happening there for various reasons. Aside from anthropogenic settlement and constant presence, construction works inside the park. The Dherua-Medinipur highway linker road also severely threatens the park’s biodiversity.

Salua range also has a significant number of avian species. Most parts are only explored once our findings are due to transportation issues that have yet to be discovered by the birding community. Equipped with sizeable semi-deciduous tree species and shrubs, the Salua range is a perfect habitat for the avian community. Most of the area still needs to be explored due to the unavailability of sighting data, inapproachable as there is no direct road link to the site areas. More detailed work is required to assess the prominent diversity data. The Dharma-Medinipur area has the main concern regarding vehicle collision and wildlife fragmentation through the construction of roads. Many local reports have been raised yearly about bird collisions with the windscreen of heavy-duty trucks and vehicles. Vadutola region has been rich in avifaunal diversity due to the dense forest of the Salboni region. Delta range has some significant biodiversity records. Overall, we assessed the parts with almost five years of biodiversity data to showcase recent trends of bird sighting and their habitat status.

## Acknowledgements

We would like to acknowledge all the volunteers who have helped us during this whole bird sighting and made this work possible.

## Contributions

N.M. and S.D. designed the whole project. Every author contributed equally during the bird survey, N.M. and S.D. wrote the manuscript, and N.M. did all the statistical analysis.

---

**Supp Fig. 1.**
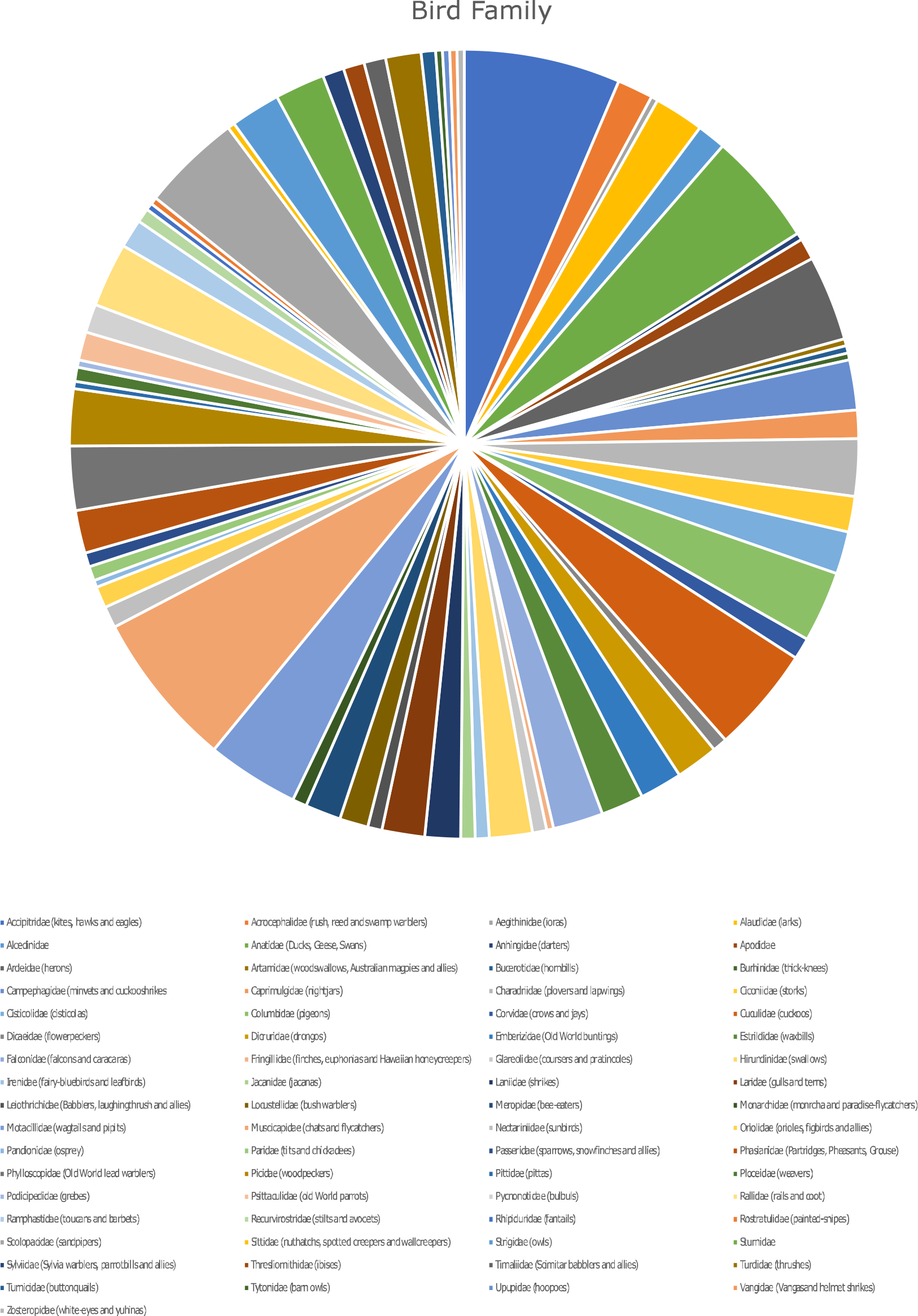
Family names of the birds.

